# A simple and effective sampler to collect undisturbed cores from tidal marshes

**DOI:** 10.1101/515825

**Authors:** Georgios Giannopoulos, Dong Y. Lee, Scott C. Neubauer, Bonnie L. Brown, Rima B. Franklin

## Abstract

Core sampling is a common procedure in wetland ecology. PVC tubes are widely used to collect soil cores; however several studies fail to describe even the most typical characteristics of their sampling auger. This work aims to fill this gap and provide a simple and standardized core sampler design. A simple and inexpensive sampler is described for field use. Its main advantages are: 1) extraction of undisturbed cores; 7.6 cm diameter and up to 100 cm depth, 2) it is light-weight, sturdy and reliable, 3) it is made from widely available PVC items and 4) it requires minimal skills to assemble. The sampler is introduced in the wetland soil, by rotation, to the desired depth; the tooth-like edge can cut and penetrate through the dense root systems. The sampler is capped with an industrial-type stopper, and pulled up. The core is held in the sampler by suction – negative pressure. A plunger is then used to slowly remove the soil – core and sub – samples are collected. Alternatively, both ends of sampler could be sealed and taken to the lab. An 8 cm long sub-sample was adequate for soil physical, biogeochemical and molecular analysis. The sampler has been tested for 2 years with more than 200 cores taken from tidal wetlands in Chesapeake Bay, Virginia, USA. A negative correlation between salinity and organic matter content at 3 – 5 and 8 – 10 cm was found. For the deeper samples (48 – 50 cm), a positive correlation between salinity and organic matter was observed. The sampler worked satisfactory and it required no maintenance besides cleaning.

## INTRODUCTION

Core sampling is a common procedure in wetland ecology, and depending on the nature of the investigation different core depths are required (Tiner, 1999). Wetland biogeochemists are interested in relatively short cores (cm-m range) to study the acrotelm and catotelm. Sedimentologists are generally interested in long cores (m-km range) to investigate the geologic history. Wetland soils are commonly classified as anaerobic soils. Due to their high water content the oxygen diffusion rate is orders of magnitude slower than well-drained agricultural soils. Wet conditions supress microbial respiration rates and as a result, significant amounts of organic matter accumulate. Thus the study of wetland C dynamics essentially requires undisturbed samples that retain their natural anaerobic conditions thus permitting direct and unbiased observations.

Collecting wetland samples using traditional methods and tools i.e. the Dutch auger, ultimately disturb and even destroy the sample (Dane and Topp, 2002). Several designs have been developed that aim to collect undisturbed soil samples for a wide range of wetland soils (DeLaune *et al.*, 2013). The most common samplers include the tube auger (AMS Inc., 2017), the Russian borer (LG. Franzén and T.L Ljung, 2009), the peat sampler (Buttler *et al.*, 1998), the Rickly sampler (Rickly Hydrological Co. Inc., 2017), the International Rice Center sampler (Savant and De Datta, 2008), the Caldwell PVC sampler (Caldwell *et al.*, 2005), and vibra/power corer systems (Finkelstein and Prins, 1981). Those specialized samplers could be hard and expensive to obtain; their sales price range from 400 USD to 3000 USD (AMS Inc., 2017). Vibra/power core samplers are typically provided as a service. Thus, financial considerations are often limiting when the costs of sediment retrieval are very high on a per-sample basis. Furthermore, several designs include moving or threaded parts that easily get clogged with debris, muddy soil and plant residues or require extended effort to retrieve the sample i.e. wrench, hand-pumps and valves, considerably slowing the field sampling process.

Wetland soils are heterogeneous consisting of water-saturated soil sediments with thick networks of plant material (e.g., roots, decaying plant tissues). In US tidal wetlands, the soil bulk density ranges from 0.06-2.53 g.cm^−3^ (Nahlik and Fennessy, 2016) with an average value of 0.35 g.cm^−3^ (Morris *et al.*, 2016), therefore, sampling undisturbed samples from such low density soils with available samplers could become challenging and time-consuming. Considering the above, the proposed sampler design is based on a thin and smooth wall construction to minimize friction and disturbance. The bottom edge of the pipe is bevelled and a saw-tooth pattern is made to cut through plant material and fibrous debris. A strong handle is added on the top edge of the sampler for tough handling. To retain the core, an air-tight stopper instead of a vacuum seal system (Savant and De Datta, 2008) is used for simplicity. This industrial type of stopper is expected to secure the core by negative-pressure while the sampler is pulled up. The proposed design was assembled at the Department of Biology, Virginia Commonwealth University and tested in tidal wetlands at Chesapeake Bay, Virginia, USA.

## MATERIAL AND METHODS

### Site

Samples for this study were collected in July 2015, and the sampler was used during a two-year sampling expedition (July 2015-August 2017) from six tidal marshes in Chesapeake Bay, Virginia USA (Table 1), varying in salinity from completely fresh (*ca.* 0 ppt) to mesohaline (*ca.* 10 ppt). Tidal marshes along the Pamunkey and York River range in size from small pocket and fringing marshes <0·5 ha to expansive marshes up to 587 ha (Baldwin *et al.*, 2012). The dominant plant species were typical of freshwater, oligohaline and mesohaline wetlands and consisted of *Peltandra virginica, Zizania aquatica, Polygonum arifolium, Sagittaria latifolia, Phragmites australis* and *Spartina cynosuroides.* Higher plant diversity is typically found at oligohaline sites, e.g. Sweet Hall Marsh, than the lower and more saline marshes, e.g. Taskinas Creek where a monoculture of *Spartina cynosuroides* was found; corroborating earlier observations of Perry and Atkinson (2009).

**Table 1.** Site geographic coordinates

| Site Index | Site | Latitude (N) | Longitude (E) |
| --- | --- | --- | --- |
| 1 | Pamunkey Reservation | 37.57115 | -77.0225 |
| 2 | White Oak Landing | 37.56811 | -76.9133 |
| 3 | Cumberland Marsh | 37.55726 | -76.9727 |
| 4 | Sweet Hall Marsh | 37.55776 | -76.8768 |
| 5 | West Point Marsh | 37.54914 | -76.8256 |
| 6 | Taskinas Creek Marsh | 37.41592 | -76.7144 |

### Sampler construction and assembly

The sampler was made exclusively with PVC materials widely available in major hardware stores. The auger consisted of a pipe, a coupling, a flange for strength and grip, and a stopper. We used a custom-made plunger to remove the soil core from the sampler. The plunger was assembled with the following PVC materials; a pipe, a cap and a disk (Figure 1; Table 2).

**Figure 1.**
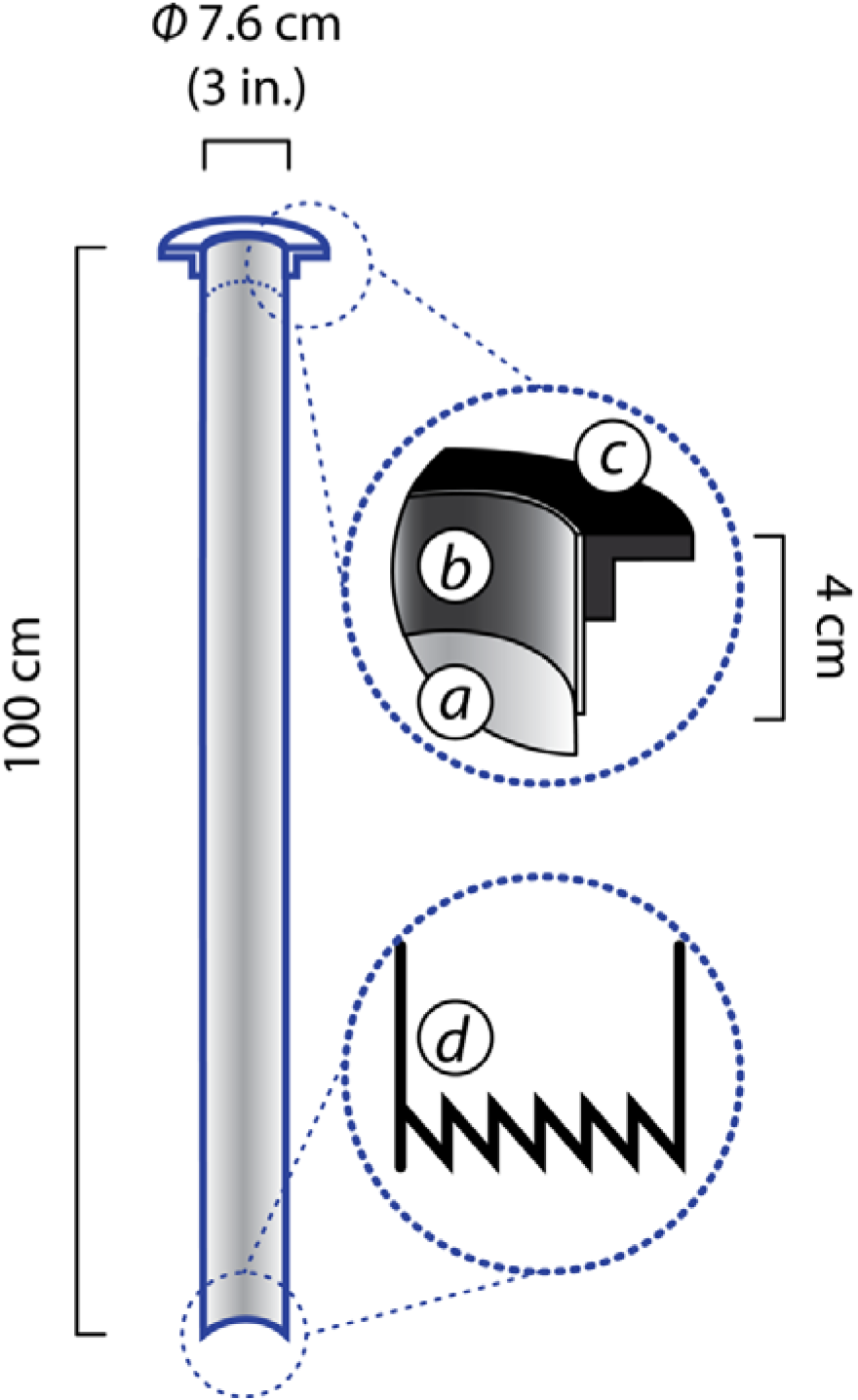
Schematic illustrating a cross-section of the sampler; a) Φ3 in. drain pipe, b) coupling, c) flange and d) detail of the saw-tooth finishing. Details about the components used are listed in Table 2

**Table 2.** Materials used to assemble the sampler and the plunger

| Item* | Product Description | Manufacturer |
| --- | --- | --- |
| PVC Sewer & Drain Pipe Φ3 in. – cut at 100 cm length | #PVC-30030 Sewer and Drain Pipe 3-in. x 10-ft. ASTM D 2729 | Charlotte Pipe, USA |
| Stopper Φ3 in. | #33402 Gripper – 75 mm, Test Plug | Oatey, USA |
| PVC Drain Fitting - Flange Φ3 in. | #43508 Closet Flange with plastic ring without test cap ASTM PVC D2665 ABS D2661 B181.1 | Oatey, USA |
| PVC Coupling Φ3 in. – cut at 4 cm length | #3P05 | NDS, USA |
| PVC pipe Φ1 in. – cut at 120 cm length | #PVC-04010-1000 1-in x 5-ft 450 Schedule 40 PVC Pipe ASTM D 178 | Charlotte Pipe, USA |
| PVC cap | #C447-010 1 in. Schedule 40 PVC Cap | DURA, USA |
| PVC disk Φ3 in. | Custom made PVC disk Φ3 in.* | N/A |
| Stainless screw with nut (M8) | M8x20 panhead screw, M8 nut with safety washer | N/A |
| PVC Cement & Primer | #302483 8 oz. PVC Cement and Purple Primer Handy Pack | Oatey, USA |

\* PVC disk was constructed by breaking off the horizontal flange around a $\Phi$ 3 in. test cap fitting (#00131-1000; Charlotte Pipe, USA). Other methods of constructing disks from PVC sheets could be used.

PVC pipes with nominal diameter (Φ) 1 in. and 3 in. and the coupling were cut to the desired length as indicated in Table 2. Few tools were required; a handsaw, a file, a screw driver, a M8 wrench and a piece of 80 grit sandpaper. Firstly, the lower edge of the pipe (Φ 3 in.) was bevelled with a piece of rough sandpaper and a saw-tooth pattern was created using a file. Secondly, PVC items were cleaned and glued with a two – component PVC cement, as follows. The sampler components were glued in order; coupling with pipe (Φ 3 in.; top-edge), then the flange with the glued pipe-coupling (Figure 1). For the plunger, the custom-made PVC disk was attached with the PVC cap (top) with a screw. Then the assembled cap-disk was glued on the top end of the Φ1 in. pipe (Figure 2).

**Figure 2.**
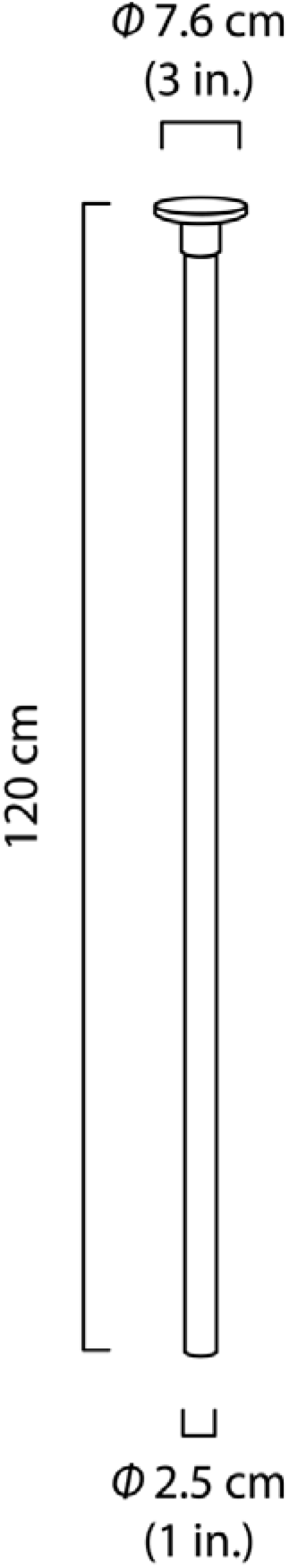
Schematic illustrating the plunger. Details about the components used are listed in Table 2.

### Core Sampling Example

Core sampling was easily carried out by one person; however two persons would speed up the process. The sampler was placed upright on the desired spot, flange on top and bevelled edge on the ground. Then the sampler was forced slowly into the wetland soil with a simultaneous rotation (Figure 3a). The rotation cut any root or plant material. Once the sampler reached the desired depth, the top was sealed air-tight with an industrial stopper (Figure 3b). Then the sampler was pulled out of the ground. If desired, the sealed sampler could be transported to the lab; in that case it is advisable to seal the bottom edge of the sampler with another stopper. The plunger was placed underneath the sampler (bevelled edge) and the stopper was removed. The sampler was pushed down, and sub-samples were sectioned off the top of the sampler (flange edge. Figure 3c). Subsamples for 3-5, 8-10 and 48-50 cm from the surface were immediately sealed in air-tight bags (Zip-loc, USA) and stored at 4°C until processing. The sampler and plunger was cleaned and reused.

**Figure 3.**
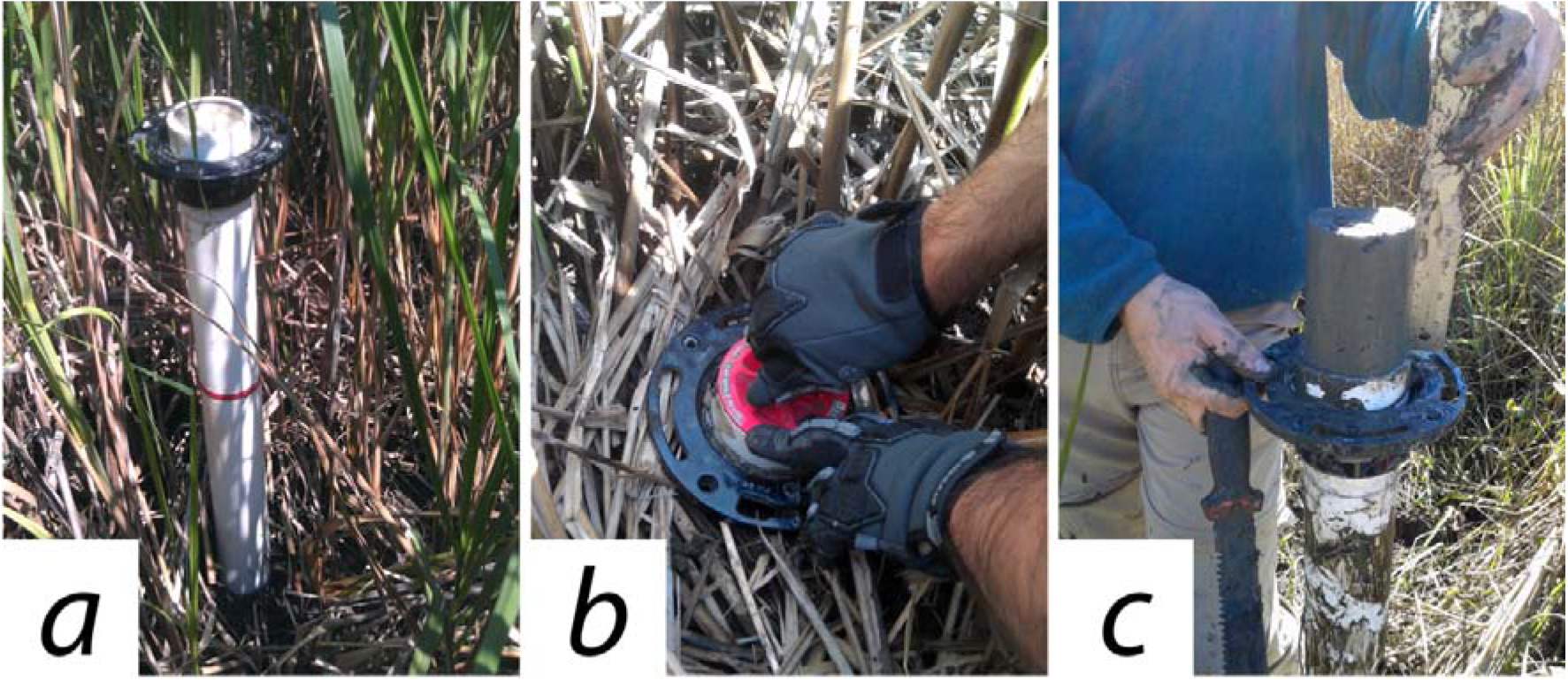
Photos during sampling: a) sampler placed at the desired spot, b) after being pushed into ground, sampler capped with an industrial-type stopper, c) sub-samples sliced off at desired depths.

### Field Testing

The performance of different internal diameter (Φ1, 2, 3 and 4) sampler was tested. The difference in length between the core acquired-and the sampler inserted-length was recorded (n=4) at the field. Notes on ease-of-use and component performance were also taken.

### Determination of water content, total organic matter and soil-water salinity

From *ca.* 90 cm^3^ sample, 15 g wet weight sediment were used to determine, firstly the water content gravimetrically (oven-dried for 72 h at 70 °C) and secondly the total organic matter (loss-on-ignition; 550 °C for 6 h) of each sample. Additionally, *ca.* 50 g wet weight sediment were used to extract soil-water by centrifugation (6000 RPM × 30 mins). The supernatant was filtered (0.2 μm Millex GP, Millipore, USA). The Cl^−^ concentration of the supernatant was determined with ion-chromatography (Dionex ICS-5000; Thermo, USA) and used to calculate salinity (Bianchi, 2007).

### Statistical evaluation

The differences in sample length, water content, carbon and salinity were statistically tested with ANOVA; Tukey-HSD in RStudio (R Core Team, 2016). All variables were log2-normalized.

## RESULTS

The undisturbed soil core sampling technique was tested successfully in tidal wetland soil, with or without flood water. Two persons could collect 15 undisturbed soil cores with several sub-samples for biogeochemical and molecular analyses in approximately four hours. The sampler weighs < 1 Kg and was assembled in approximately 1 h. The materials cost approximately 45 USD plus sales tax (Lowes, Virginia, USA; July 2015). The sampler was used to collect > 200 cores during a two-year sampling expedition (July 2015-August 2017) from six tidal marshes in Chesapeake Bay, Virginia USA. A subset (August 2015; 54 data points) of the larger dataset is presented here. At the end of the sampling expedition, the sampler didn’t show any visual signs of damage or wear. The wing-nut from the stopper and the Φ3 plunger disk had to be replaced once, because one of the wings broke from over-tightening and the disk cracked. A thicker PVC disk could be more resilient to cracking. No other replacement was required. When working with two persons, a piece of rope was attached through the screw-holes of the flange to facilitate the extraction of the sampler. Care should be taken to force the sampler into the wetland soil in a steady and slow fashion to avoid accidental compaction. Additionally, when working in e.g. *Spartina alterniflora* dominated marshes, a long machete was carefully pushed down by the outer side of the sampler to cut roots/rhizomes and assist the process.

Four samplers with different nominal diameter (Φ) were tested; 1 in., 2 in., 3 in. and 4 in. The Φ 1 in. pipe failed to collect a 100 cm sample; while being forced in the wetland soil it became quickly clogged and only 15.3 cm (SE ±4.2, n=4) were collected. The Φ2, Φ3 and Φ4 in. pipe collected deep samples ~100 cm. There wasn’t any significant difference in sample length retrieved among the Φ2, Φ3 and Φ4 in. pipes (*p* >0.05; Figure 4). However, the volume of the sample enclosed in the Φ2 in. pipe wasn’t enough for the required lab analysis. Also, the Φ4 in. pipe, when filled with a 100 cm core, was very heavy for a single person to pull up from the ground (drag-force). Therefore the sampler design was based on a Φ3 pipe, a trade-off between collected volume and ease of use.

**Figure 4.**
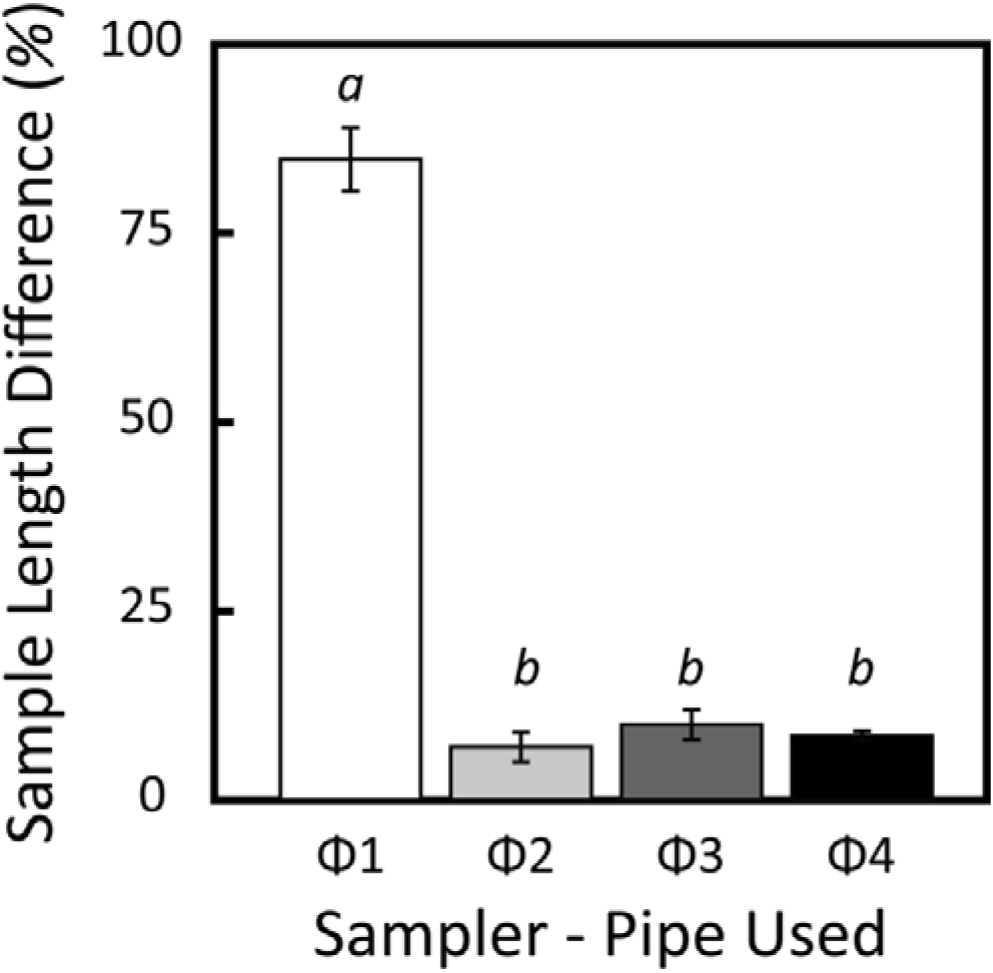
Average sample length difference (± SD, n = 4) retrieved from a 100 cm long pipe. Treatments sharing the same letter are not significantly different (Tukey HSD, *α*=0.05).

The water content of the samples was on average 70% (SE±2) and the organic matter content was 22 % (dw, SE±2). There was a negative correlation between salinity and organic matter content at 3-5 and 8-10 cm below the surface (Figure 5). For the deeper samples (48-50 cm), reaching a gleyed clay horizon, a positive correlation between salinity and organic matter was found.

**Figure 5.**
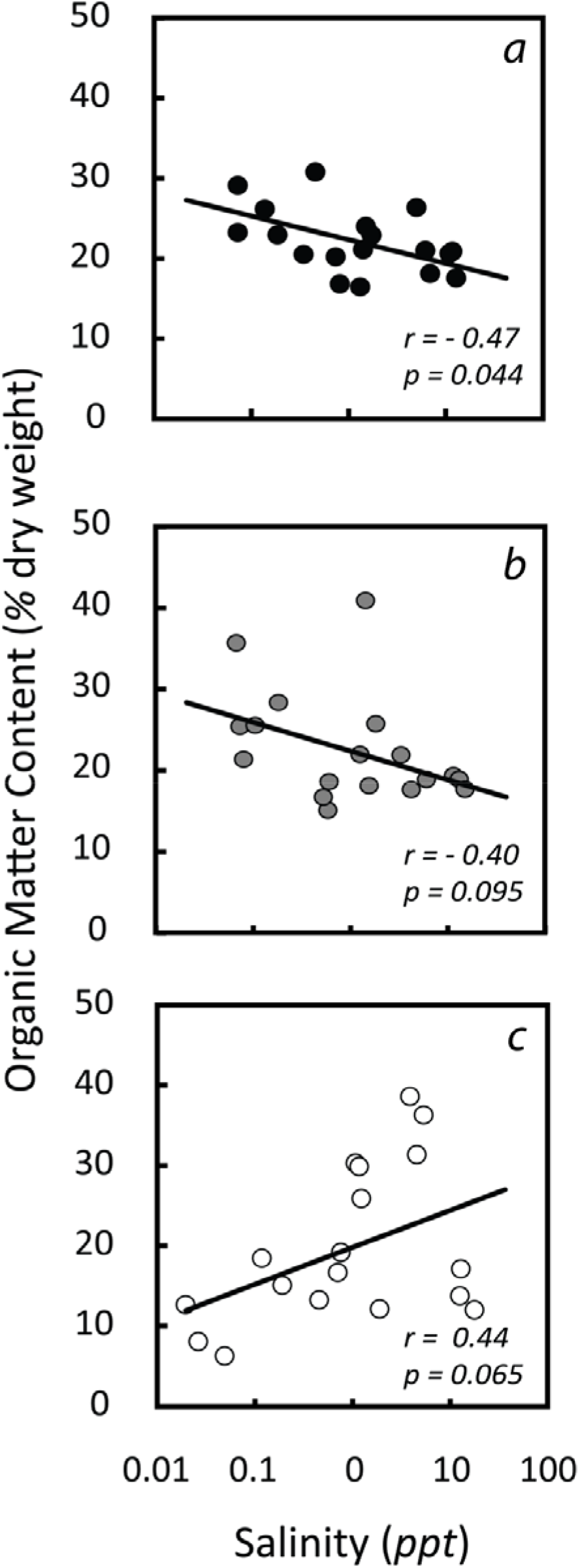
Pearson correlations of organic matter content (% DW) with pore-water salinity at three sampling depths: a) 3 – 5, b) 8 – 10 and c) 48 – 50 cm from the surface.

## DISCUSSION

Several soil samplers have been developed for collecting undisturbed cores. Their commercial availability is limited and their cost is frequently expensive. Their designs aim to collect undisturbed cores; however complicated designs with various moving parts can make the actual procedure of core-sampling rather cumbersome. In this study, a simple, low-cost, light-weight, sturdy and effective sampler was designed and tested in tidal wetland soils of Chesapeake Bay. The sampler is made with non-corrosive materials that minimize any chemical interference (e.g. iron rust) in the sample. It was tested for 2 years with more than 200 cores taken from tidal wetlands in Chesapeake Bay, Virginia, USA. A relationship between salinity and organic matter content was found, which is typical of tidal wetlands (Morrissey *et al.*, 2014a). It worked satisfactorily and it required no maintenance besides cleaning.

The physical limitations of the sampler depend on the method of sampler advancement and the nature of the soil matrix. This sampler was tested in wetland organic soils and was pushed into the soil manually. Due to the nature of the materials used (PVC), high-impact hammering is not recommended. The maximum tested insertion length was 100 cm. The propose design could be extended to greater length; however additional tests should be done prior to routine field work. Several researchers add water prior to sealing the sampler to saturate the sample and generate an air-free seal (Caldwell *et al.*, 2005, Finkelstein and Prins, 1981). Such technique could also be applied to the proposed sampler design if required. However, the addition of water could impact the pore-water chemistry; therefore it was avoided for this study.

The sampler is particularly appropriate for regular wetland sampling for research or monitoring purposes. Furthermore, this low cost sampler could be intergraded in field workshops and training activities. The use of PVC sampling devices is becoming popular because of their light weight and ease-of-use in remote areas; the most common designs are made with a single PVC tube cut at the desired length. For example Livolsi *et al.* (2014) used a PVC sampler to assess seed and invertebrate feed in wetlands (diameter: 5.1 cm; length: 12.7 cm). Other simple sampling designs include for example a modified syringe barrel (60 ml) to collect soil samples for microbial analysis (Morrissey *et al.*, 2014b). This sampler collects 45 ml (cm^3^) per cm depth which is ample for molecular determinations. Thus, besides the evaluation of physical and chemical properties of wetland soils, this sampler could also be used to collect samples for the study of root morphology, sedimentation and soil microbiological aspects.

Although PVC tubes are widely used to collect soil cores, several studies fail to describe even the most typical characteristics of their sampling auger (e.g. diameter) or construction methods. This work aims to fill this gap and provide a simple and standardized core sampler design. In conclusion, the proposed design was effective in collecting undisturbed samples from wetland soil, sturdy in operation and low-cost to construct. The core-samples could be sub-sampled at the field or at the laboratory for further analysis.

## AKNOWLEDGEMENTS

This work was funded by a National Science Foundation grant to Rima Franklin, Bonnie Brown, and Scott Neubauer (NSF-DEB-1355059, “Climate change effects on coastal wetlands-Linking microbial community composition and ecosystem responses”).

## LITERATURE CITED

AMS INC. 2017. AMS Sampling Tools [Online], https://www.ams-samplers.com/. Available: https://www.anns-samplers.com/.

Baldwin, A. H., Kangas, P. J., Megonigal, J. P., Perry, M. C., Whigham, D. F. & Batzer, D. P. 2012. Coastal Wetlands of Chesapeake Bay. In: darold P. batzer & Andrew H. Baldwin (eds.) Wetland habitats of North America: Ecology and conservation concerns. Berkeley CA, USA: University of California Press.

Bianchi,T. S. 2007. Biogeochemistry of Estuaries, New York, USA, Oxford University Press.

Buttler, A., Grosvernier, P. & Matthey, Y. 1998. A new sampler for extracting undisturbed surface peat cores for growth pot experiments. New Phytologist, 140,355–360.

Caldwell, P. V., Adams, A. A., Niewoehner, C. P., Vepraskas, M. J. & Gregory, J. D. 2005. Sampling Device to Extract Intact Cores in Saturated Organic So’ús. Soil Science Society of America Journal, 69, 2071.

Dane J. H. &Topp, C. G. 2002. Methods of Soil Analysis: Part 4 Physical Methods, Madison, WI, Soil Science Society of America.

Delaune, R. D., Reddy, K. R., Richardson, C. J. & Megonigal, J. P. 2013. Methods in Biogeochemistry of Wetlands, Madison, Wl, Soil Science Society of America.

Finkelstein K. & Prins D. 1981. An Inexpensive, Portable Vibracoring System for Shallow-Water and Land Application. Coastal Engineering Technical Aid. Coastal Engineering Research Center.

L.G. Franzén & T.L. Ljung 2009. A carbon fibre composite (CFC) Byelorussian peat corer. Mires and Peat, 5,1–9.

Livolsi, M. C., Ringelman, K. M. & Williams, C. K. 2014. Subsampling Reduces Sorting Effort for Waterfowl Foods in Salt-Marsh Core Samples. Journal of Fish and Wildlife Management, 5, 380–386.

Morris J. T., Barber D. C, Callaway J. C, Chambers R., Hagen S. C., Hopkinson C. S., Johnson B. J.,Megonigal, P., Neubauer, S. C., Troxler, T. &Wigand,C. 2016. Contributions of organic and inorganic matterto sediment volume and accretion in tidal wetlands at steady state. Earths Future, 4,110–121.

Morrissey, E. M., Gillespie J. L, Morina, J. C. & Franklin, R. B. 2014a. Salinity affects microbial activity and soil organic matter content in tidal wetlands. Global Change Biology, 20,1351–1362.

Morrissey, E. M., Gillespie J. L, Morina, J.C. & Franklin, R. B. 2014b. Salinity affects microbial activity and soil organic matter content in tidal wetlands. Glob Chang Biol, 20, 1351–62.

Nahlik A. M. & Fennessy M. S. 2016. Carbon storage in US wetlands. NatCommun, 7, 13835.

Perry J. E. & Atkinson R. B. 2009. York River Tidal Marshes. Journal of Coastal Research, 10057, 409.

R Core Team 2016. R: A language and environment for statistical computing. Vienna, Austria

RICKLY HYDROLOGICAL CO. INC. 2017. Sediment Sampling [Online]. Available: http://ricklv.com/sediment-sampline/ [Accessed 07 Dec 2017.

Savant N. K. & De Datta S. K. 2008. An undisturbed core sampling and sectioning technique for wetland rice soils. Communications in Soil Science and Plant Analysis, 10,775–783.

Tiner R. 1999. Wetland Indicators, Boca Raton, CRC PRess.

